# Effects of partial selfing on the equilibrium genetic variance, mutation load and inbreeding depression under stabilizing selection

**DOI:** 10.1101/180000

**Authors:** Diala Abu Awad, Denis Roze

## Abstract

This preprint has been reviewed and recommended by Peer Community In Evolutionary Biology (http://dx.doi.org/10.24072/pci.evolbiol.100041).

The mating system of a species is expected to have important effects on its genetic diversity. In this paper, we explore the effects of partial selfing on the equilibrium genetic variance *V_g_*, mutation load *L* and inbreeding depression *δ* under stabilizing selection acting on a arbitrary number *n* of quantitative traits coded by biallelic loci with additive effects. Overall, our model predicts a decrease in the equilibrium genetic variance with increasing selfing rates; however, the relationship between self-fertilization and the variables of interest depends on the strength of associations between loci, and three different regimes are observed. When the *U*/*n* ratio is low (where *U* is the total haploid mutation rate on selected traits) and effective recombination rates are sufficiently high, genetic associations between loci are negligible and the genetic variance, mutation load and inbreeding depression are well predicted by approximations based on single-locus models. For higher values of *U*/*n* and/or lower effective recombination, moderate genetic associations generated by epistasis tend to increase *V_g_*, *L* and *δ*, this regime being well predicted by approximations including the effects of pairwise associations between loci. For yet higher values of *U*/*n* and/or lower effective recombination, a different regime is reached under which the maintenance of coadapted gene complexes reduces *V_g_*, *L* and *δ*. Simulations indicate that the values of *V_g_*, *L* and *δ* are little affected by assumptions regarding the number of possible alleles per locus.

## INTRODUCTION

Genetic diversity maintained within populations plays an important role in defining their adaptive potential (for a species to evolve, there must be heritable phenotypic variation on which selection can act). The ultimate source of this diversity is mutation, with a substantial proportion of new mutations being of a slightly deleterious nature (Eyre-Walker and Keightley, 2007): hence, a corollary to the maintenance of genetic diversity is the existence of a mutation load, defined as the reduction in mean fitness of a population relative to the fitness of an optimal genotype (Haldane, 1937). Furthermore, the fact that most deleterious alleles are partially recessive causes inbred offspring to have a lower fitness (on average) than outbred ones, as they tend to carry higher numbers of homozygous mutations (inbreeding depression, Charlesworth and Charlesworth, 1987).

By affecting the average degree of homozygosity of individuals and the efficiency of recombination between loci, the reproductive system of a species is expected to have an important influence on the effect of selection against deleterious alleles, and thus on the mutation load, inbreeding depression and level of diversity maintained within populations. One mating system that has received considerable attention is self-fertilization, a reproductive strategy occurring at various rates in an important proportion of plant and animal species (Jarne and Auld, 2006; Goodwillie et al., 2005; Igic and Kohn, 2006). Self-fertilization, and inbreeding in general, may have different effects on genetic polymorphisms depending on the strength of selection acting on them (Gl´emin, 2007). When directional selection against deleterious alleles is sufficiently strong relative to drift (*N_e_s* ≫ 1), the increased homozygosity caused by inbreeding is expected to improve the efficiency of selection against those alleles (purging), reducing the mutation load and inbreeding depression (Lande and Schemske, 1985; Charlesworth et al., 1990). At the other extreme, polymorphism at neutrally-behaving loci (*N_e_s* ≪ 1) should also be lowered by inbreeding, as the effective population size is reduced by identity-by-descent within loci (Pollak, 1987) and by stronger interference effects between loci — background selection, hitchhiking (Nordborg, 1997; Gl´emin and Ronfort, 2013; Roze, 2016). In intermediate regimes (*N_e_s* ∼ 1), however, the reduction in *N_e_* due to inbreeding may cause an increased frequency of deleterious alleles (because selection is less effective), which may explain the higher *π_N_ /π_S_* ratio observed in various selfing species compared with their outcrossing relatives (Brandvain et al., 2013; Burgarella et al., 2015, and other references listed in Table 1 of Hartfield, 2015).

**Table 1:**
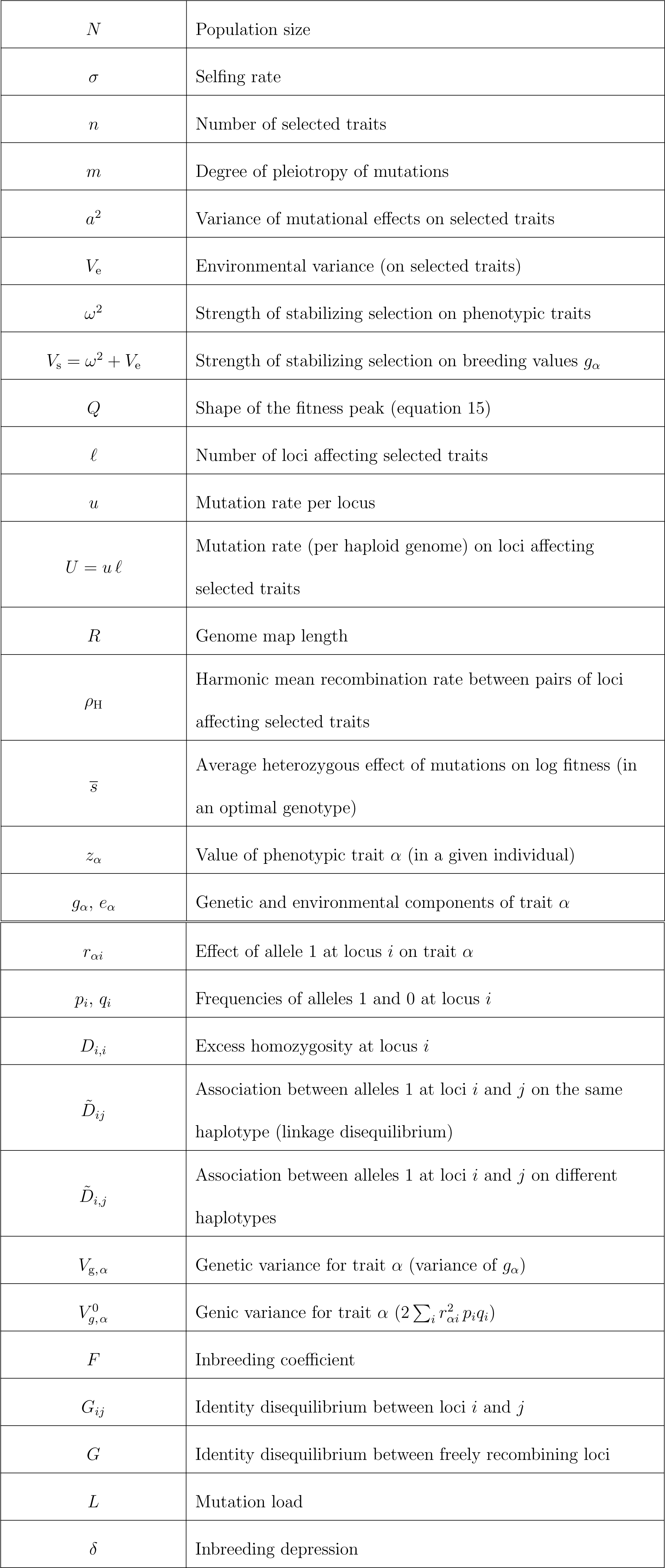
Parameters and variables of the model.

Most classical results on the effects of selfing on genetic diversity, mutation load and inbreeding depression are based on single-locus models, and thus neglect the effects of linkage disequilibria and other forms of genetic associations among loci. Previous analytical and simulation models showed that intermediate selfing rates generate correlations in homozygosity between loci, termed “identity disequilibria” (Weir and Cockerham, 1973; Vitalis and Couvet, 2001), which tend to reduce the efficiency of purging when deleterious alleles are partially or fully recessive (an effect called “selective interference” by Lande et al., 1994). When the number of highly recessive mutations segregating within genomes is sufficiently high, these correlations in homozygosity may entirely suppress purging unless the selfing rate exceeds a given threshold (Lande et al., 1994; Scofield and Schultz, 2006; Kelly, 2007; Roze, 2015). Linkage disequilibrium corresponds to another form of association between loci that may also affect the efficiency of selection: in particular, selection may be strongly limited by Hill-Robertson effects in highly selfing populations, due to the fact that selfing reduces the efficiency of recombination between loci — recombination having no effect when it occurs in homozygous individuals (Kamran-Disfani and Agrawal, 2014; Hartfield and Gl´emin, 2016). Epistatic interactions represent another possible source of linkage disequilibrium between selected loci. Charlesworth et al. (1991) considered a model in which epistasis between deleterious alleles is fixed and synergistic (the effects of mutations alone being smaller than when combined with others), and showed that the effect of the selfing rate on the load and inbreeding depression may be non-monotonic under this form of epistasis, with an increase in both variables above a (high) self-fertilization threshold. However, although models with fixed epistasis have lead to important insights, epistatic interactions are known to vary across pairs of loci, and this variation may have important evolutionary consequences (Phillips et al., 2000; Martin et al., 2007). Interestingly, several aspects of the complexity of epistatic interactions (such as possible compensatory effects between deleterious alleles, *i.e.*, reciprocal sign epistasis) are captured by models of stabilizing selection acting on quantitative traits, such as Fisher’s geometric model (Fisher, 1930). Furthermore, the distributions of epistasis generated by this type of model seem compatible with our empirical knowledge on epistasis (Martin et al., 2007).

Only a few models have explored the effect of self-fertilization on genetic variance for quantitative traits at equilibrium between mutation and stabilizing selection. Modeling a quantitative trait coded by additive loci, Wright (1951) showed that, in the absence of selection, the genetic variance for the trait is increased by a factor 1 + *F* (where *F* is the inbreeding coefficient), due to the increased homozygosity of the underlying loci. Selection will oppose this increase in variance, however, by eliminating genotypes that are too far from the optimum (purging). Stabilizing selection is also known to generate positive linkage disequilibrium between alleles at different loci having opposite effects on the trait (Bulmer, 1971), the immediate consequence of which is to reduce the genetic variance. These linkage disequilibria should also affect the efficiency of selection at each locus, and thus indirectly affect the genetic variance. Lande (1977) proposed a model of stabilizing selection acting on a single trait coded by additive loci in a partially selfing population, in which a Gaussian distribution of allelic effects is assumed to be maintained at each locus. He found that, as the increase in variance due to homozygosity is exactly compensated by the effect of purging, and the decrease in variance caused by linkage disequilibria is exactly compensated by the decreased efficiency of selection acting at each locus due to these linkage disequilibria, overall the equilibrium genetic variance is not affected by the selfing rate of the population. More recently, Lande and Porcher (2015) extended this model to multiple selected traits, and used a method developed by Kelly (2007) to take into account the effects of correlations in homozygosity across loci by splitting the population into selfing age classes (corresponding to classes of individuals having the same history of inbreeding), while assuming a Gaussian distribution of allelic effects at each locus within each class. Numerical iterations of the model showed that above a threshold selfing rate, a different regime is reached, in which strong compensatory associations between alleles at different loci reduce the genetic variance and may generate outbreeding depression (*i.e.*, lower fitness of outcrossed offspring relative to selfed offspring).

The hypothesis made by Lande (1977) and Lande and Porcher (2015) of a Gaussian distribution of allelic effects maintained at each locus (either in the whole population or in each selfing age class) has been criticized on the grounds that it implicitly assumes an unrealistically high mutation rate per locus and/or very weak fitness effects of mutations (Turelli, 1984). Lande and Porcher (2015) also considered an infinitesimal model (in which traits are coded by an infinite number of loci, selection having a negligible effect on allele frequencies at each locus), and showed that a similar threshold pattern emerges, although the effect of selfing on the genetic variance and inbreeding depression above the threshold differs between the two models (in particular, outbreeding depression is not observed in the infinitesimal model). However, the effect of selfing on the genetic variance of quantitative traits under more general assumptions regarding the strength of selection at the underlying loci remains unclear. In this paper, we introduce partial self-fertilization into previous models of stabilizing selection acting on quantitative traits coded by biallelic loci (Latter, 1960; Bulmer, 1972; Barton, 1986, 1989; Turelli and Barton, 1990; Roze and Blanckaert, 2014). Charlesworth and Charlesworth (1995) had extended such models to take complete selfing into account, and found that the genetic variance under stabilizing selection should be lower under complete selfing than under random mating. The present paper generalizes these results to arbitrary selfing rates and multiple selected traits. Assuming additive effects of alleles on phenotypes (no dominance or epistasis on phenotypic traits), we develop approximations incorporating the effects of pairwise associations between selected loci, and compare these approximations with results from individual-based simulations. Our results indicate that different regimes are possible depending on the effect of genetic associations on the genetic variance, this effect increasing as the overall mutation rate *U* and selfing rate *σ* increase, and decreasing as the genome map length *R*, mean fitness effect of mutations *s̅* and number of selected traits *n* increase. When *U* and *σ* are sufficiently low and *R*, *s̅* and *n* sufficiently high, the effect of associations is negligible and the mutation load, inbreeding depression and genetic variance are well predicted by classical expressions ignoring associations. As the strength of associations increases, a second regime is entered in which the overall effect of associations is to reduce purging, thereby increasing the genetic variance, mutation load and inbreeding depression; this “weak association” regime is generally well predicted by our approximations which include the effects of pairwise associations between loci. For yet higher *U*, *σ* and/or lower *R*, *s̅* or *n*, a third regime is reached in which strong associations between loci caused by compensatory effects among mutations reduce the genetic variance, load and inbreeding depression. Although our approximations break down in this “strong association” regime, the approximation proposed by Charlesworth and Charlesworth (1995) provides accurate results under complete selfing when the mutation rate is sufficiently high and the mean fitness effect of mutations sufficiently low.

## MODEL

### Genotype-phenotype map

The parameters and variables of our model are summarized in Table 1. We consider a diploid population of size *N* with discrete generations. Offspring are produced by self-fertilization with probability *σ*, and by random union of gametes with probability 1 *− σ*. The fitness of an organism represents its overall relative fecundity (assumed very large for all individuals), and depends on the values of *n* quantitative phenotypic traits under stabilizing selection. In the following we use subscripts *α*, *β*, *γ*… to denote phenotypic traits, while subscripts *i*, *j*, *k*… denote loci. The value of trait *α* in a given individual is denoted *z_α_*, and can be decomposed into a genetic and an environmental component:

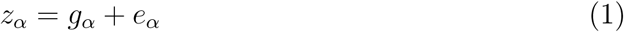

where the environmental component *e_α_* is sampled from a Gaussian distribution with mean zero and variance *V*_e_ (the same for all traits). The genetic component *g_α_* (“breeding value”) is controlled by a large number of biallelic loci with additive effects. The two alleles at each locus are denoted 0 and 1, while 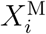 and 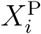 are defined as indicator variables that equal zero if the individual carries allele 0 at locus *i* on its maternally 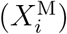 or paternally 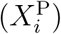 inherited chromosome, while they equal 1 if allele 1 is present. We also assume that *g_α_* = 0 in an individual homozygous for allele 0 at all loci, so that:

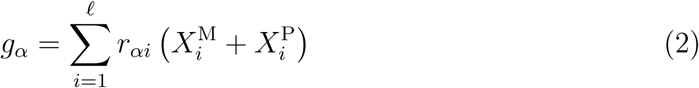

where *ℓ* is the number of loci affecting phenotypic traits, and *r_αi_* the effect on phenotype *α* of changing the allelic state of one gene copy at locus *i* from 0 to 1 (note that *r_αi_* may be negative).

Following Chevin et al. (2010), Lourenço et al. (2011) and Roze and Blanckaert (2014), a parameter *m* measures the degree of pleiotropy of mutations: each locus affects a subset of *m* phenotypic traits, sampled randomly (and independently for each locus) among the *n* traits. Therefore, *m* = 1 means that each locus affects a single trait, while *m = n* corresponds to full pleiotropy (each locus affecting all traits), as in Fisher’s geometric model (Fisher, 1930). We assume that the distribution of mutational effects *r_αi_* over all loci affecting trait *α* has average zero and variance *a*^2^ (the same for all traits); if locus *i* does not affect trait *α*, then *r_αi_* = 0. For simplicity, we consider a fully isotropic model with no mutational covariance between traits. Finally, *u* denotes the mutation rate from allele 0 to allele 1 and from allele 1 to allele 0 at each locus, while *U* = *uℓ* is the haploid mutation rate over all loci (per generation).

From the previous definitions, and assuming that population size is sufficiently large, mean trait values are given by:

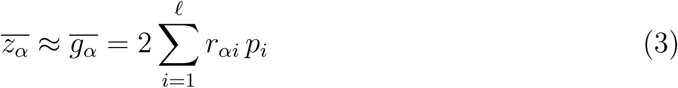

where *p_i_* is the frequency of allele 1 at locus *i*. As we assume no *G × E* interaction, the variance in trait *α* is given by:

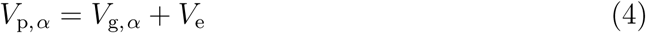

where *V*_*g, α*_ is the variance in *g_α_* (genetic variance). In the next subsection, we show how *V*_g*, α*_ can be expressed in terms of genetic associations within and between loci.

### Genetic associations and decomposition of the genetic variance

Genetic associations are defined as in Kirkpatrick et al. (2002). In particular, the centered variables *ζ_i_*_∅_ and *ζ*_∅_,*_i_* are defined as:

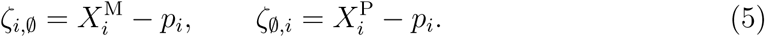

Furthermore, products of *ζ_i,∅_*, *ζ_∅,i_* variables are denoted:

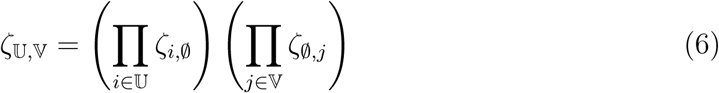

where 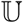 and 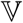 represent sets of loci. For example, for 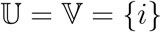, we have:

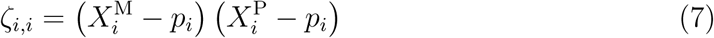

while for 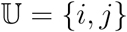 and 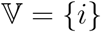:

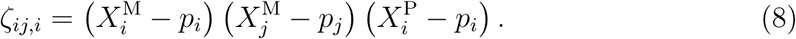

Finally, genetic associations 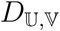 are defined as averages of 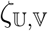 variables over all individuals:

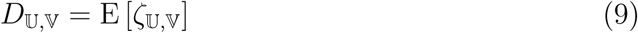

where E stands for the average over all individuals in the population. We also define 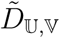 as 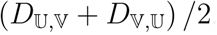, and write 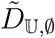 as 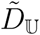 (for simplicity). In particular, *D_i,i_* is a measure of excess homozygosity (due, for example, to non-random mating) at locus *i* (*D_i,i_* = 0 at Hardy-Weinberg equilibrium). As shown in Supplementary File S1, it can be written as 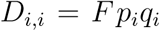, where *F* is the inbreeding coefficient (probability of identity by descent between two alleles present at the same locus in the same individual). The association *D*̃*_ij_* corresponds to the linkage disequilibrium between loci *i* and *j* (association between alleles present on the same haplotype), while *D*̃*_i,j_* is the association between alleles at loci *i* and *j* present on different haplotypes of the same individual. We will see that the association *D_ij,ij_* also appears in the computations, and can be expressed as *ϕ_ij_p_i_q_i_p_j_ q_j_*, where *ϕ_ij_* is the probability of joint identity be descent at loci *i* and *j*. The quantities *ϕ_ij_* and *F* enter into the definition of the identity disequilibrium between loci *i* and *j*, given by *G_ij_* = *ϕ_ij_* − *F*^2^ (Weir and Cockerham, 1973), which will appear in some of our results.

From these definitions, and using equations 2 and 3, the genetic variance for trait *α* can be written as:

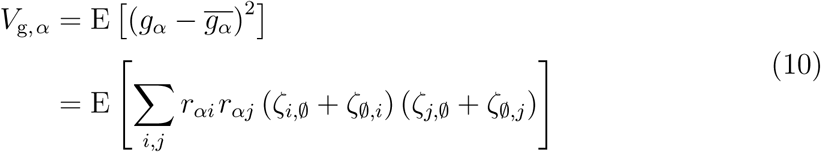

where the last sum is over all *i* and *j* (including *i = j*). Using the fact that *D*̃*_ii_* = *p_i_q_i_*, one obtains from equation 9:

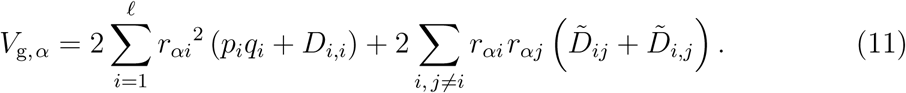

Following previous usage (e.g., Bulmer, 1985), we will call *genic variance* (denoted 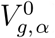) the quantity 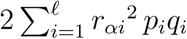, corresponding to the genetic variance in a population with the same allele frequencies, but in the absence of genetic association (within and between loci). As shown by equation 11, excess homozygosity tends to increase the genetic variance through the term in *D_i,i_*. The second term of equation 11 (the effect of between-locus associations) tends to be negative under stabilizing selection, since the allele increasing the value of trait *α* at locus *i* tends to be associated with the allele decreasing its value at locus *j* (e.g., Bulmer, 1971, 1974; Lande, 1976; Turelli and Barton, 1990). However, below we show that that excess homozygosity and associations between loci also affect equilibrium allele frequencies, and thus the genic variance.

### Fitness function

Most of the results derived in this paper assume an isotropic, Gaussian fitness function, the fitness of an individual being given by:

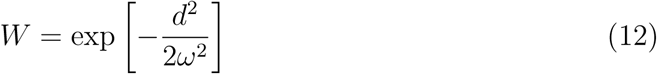

where *ω*^2^ measures the strength of selection, and 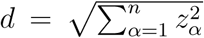 is the Euclidean distance (in phenotypic space) between the individual’s phenotype and the optimum, which we assume is located at **z** = (0, 0,…, 0). From equation 12, the fitness associated with a given genotype (obtained by averaging over environmental effects) is also Gaussian, and given by:

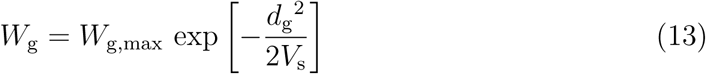

with *V_s_* = *ω*^2^ + *V*_e_, *W*_g,max_ = (*ω*^2^/*V_s_*)^*n*/2^ (the mean fitness of an optimal genotype), and 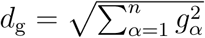 (the Euclidean distance between the breeding value of the individual and the optimum). Under our mutational model, the mean reduction in log *W*_g_ caused by a heterozygous mutation present in an optimal genotype is:

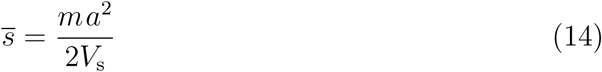

(e.g., Martin and Lenormand, 2006b). Under our assumption of additivity of phenotypic effects it is easy to show that the reduction in log *W*_g_ caused by a homozygous deleterious allele (in an optimal genotype) is four times the reduction caused by the same allele in the heterozygous state. Provided that most mutations have weak fitness effects (so that log (1 − *s*) ≈ −*s*), the dominance coefficient of deleterious alleles is thus close to 0.25 at the fitness optimum (see Manna et al., 2011 for more general results on dominance in Fisher’s geometric model).

The effect of the shape of the fitness peak will be explored using a generalized version of equation 13 (e.g., Martin and Lenormand, 2006a; Tenaillon et al., 2007):

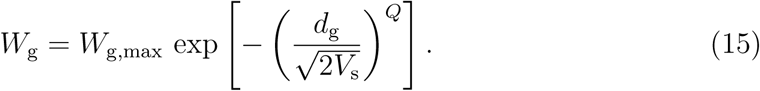

Gaussian fitness (equation 13) thus corresponds to *Q* = 2, while the fitness peak is sharper around the optimum when *Q* < 2, and flatter when *Q* > 2. Importantly, *Q* affects the average dominance coefficient of deleterious alleles, making them more dominant for *Q* < 2 and more recessive for *Q* > 2 (Manna et al., 2011), as well as the average epistasis (on fitness) between alleles, positive for *Q* < 2, and negative for *Q* > 2 (Gros et al., 2009). Approximations for the mutation load and inbreeding depression can be derived for *Q* ≠ 2 as long as the distribution of breeding values in the population is approximately Gaussian.

### Individual-based simulations

In order to verify the analytical results obtained, individual-based simulations were run using a C++ program described in Supplementary File S5 (and available from Dryad), in which the genome of each individual consists of two copies of a linear chromosome carrying *ℓ* equidistant biallelic loci affecting the *n* traits under selection. Another version of the program was used to consider a different genetic architecture, under which an infinite number of alleles are possible at each locus (see Supplementary File S5).

## RESULTS

### Neglecting associations between loci

In the following section we show that genetic associations between loci may be neglected when the haploid genomic mutation rate *U* is sufficiently low. In this case, equation 11 simplifies to:

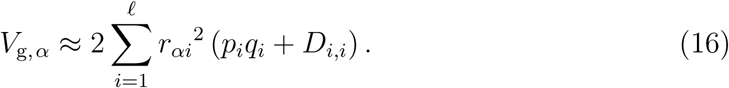

Expressions for *p_i_q_i_* and *D_i,i_* at equilibrium, assuming weak selection (*V*_g*, α*_ *≪ V*_s_) and neglecting associations among loci are derived in Supplementary File S1. To leading order, 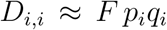 where *F* = σ/(2 − *σ*) is the inbreeding coefficient. Neglecting associations between loci and assuming that mean phenotypes are at the optimum (*g_α_* = 0), the effect of selection on *p_i_q_i_* is given by:

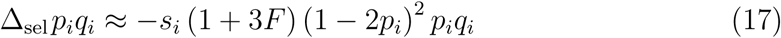

where 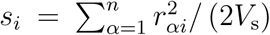 is the heterozygous effect of a mutation at locus *i* on log fitness in an optimal genotype. Furthermore, because mutation changes *p_i_* to *p_i_* (1 *− u*) + *u* (1 *− p_i_*), the change in *p_i_q_i_* due to mutation is (to the first order in *u*):

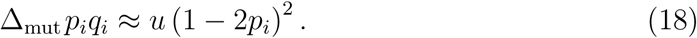

In regimes where genetic drift can be neglected, Δ_sel_*p_i_q_i_* = −Δ_mut_*p_i_q_i_* at mutation-selection balance, leading to either *p_i_* = 1/2 or:

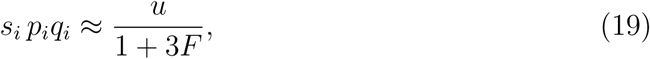

in agreement with results of previous biallelic models under random mating (e.g., Bulmer, 1972; Barton, 1986). A stability analysis indicates that the equilibrium given by equation 19 is stable when *p_i_q_i_* < 1/4 (that is, when *s_i_* (1 + 3*F*) > 4*u*), otherwise *p_i_* = 1/2 is stable. When all loci are at the equilibrium where *p_i_q_i_* < 1/4, summing both sides of equation 19 over *i* yields, using 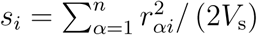:

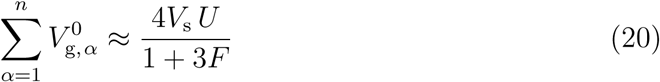

where again 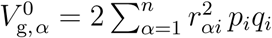 is the genic variance. By symmetry the equilibrium genic variance should be the same for all traits, and thus:

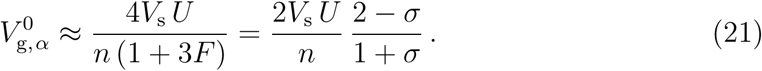

From equations 16 and 21, the equilibrium genetic variance is:

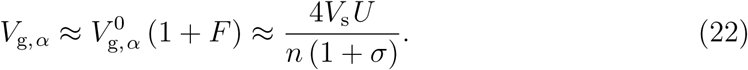

When *σ* = 0 and *n* = 1, equation 22 is equivalent to the result of previous biallelic models (e.g., Latter, 1960; Bulmer, 1972) and to Turelli’s house-of-cards approximation (Turelli, 1984).

Assuming that the variance in log-fitness is small, mean fitness is approximately 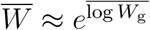. Defining the mutation load *L* as the reduction in *W̅* relative to the average fitness of an optimal genotype, one obtains from equation 13:

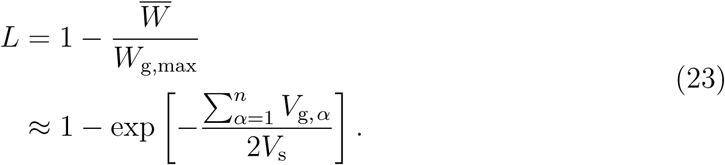

Equations 22 and 23 yield:

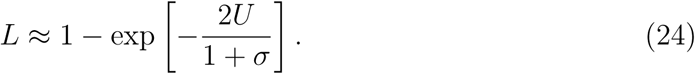

Inbreeding depression *δ* measures the mean fitness of selfed offspring, relative to the mean fitness of outcrossed offspring. Under the same assumptions, it is given by:

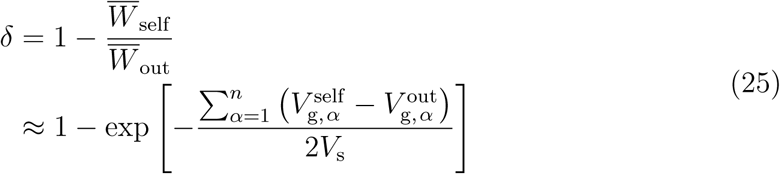

where 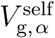 and 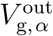 are the genetic variances for trait *α* among selfed and out-crossed offspring, respectively (e.g., Lande and Schemske, 1985). The intralocus association *D_i,i_* among selfed offspring is 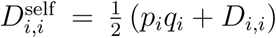 and therefore 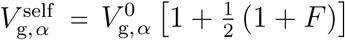, while 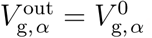, yielding (using equation 21):

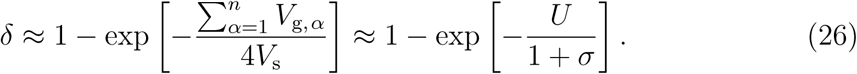

Equations 24 and 26 are equivalent to the classical expressions obtained for the load and inbreeding depression at mutation-selection balance when the dominance coefficient *h* of deleterious alleles is set to 0.25 (e.g., Charlesworth and Charlesworth, 1987), in agreement with the fact that *h ≈* 0.25 under Gaussian stabilizing selection when mutations have additive effects on phenotypes (see previous section).

Figure 1 shows that the mutation load is well predicted by equation 24 when *N_e_s̅* is sufficiently large (for *U* = 0.1 and *n* = 50), and generally decreases as selfing increases — results for different numbers of loci *ℓ* are shown in Supplementary Figure S1, while Supplementary Figures S2 and S3 show that the genetic variance and inbreeding depression follow similar patterns. Drift may have significant effects on genetic variation, however, when *N_e_s̅* is ≈ 1 or lower. Following Bulmer (1972), a diffusion model can be used to compute the expected value of *p_i_q_i_* under selection, mutation and drift, provided that the effects of associations between loci are neglected. As explained in Supplementary File S2, the result can then be integrated over the distribution of *s_i_* across loci to obtain the equilibrium genetic variance, inbreeding depression and mutation load. Figures 1 and S1 – S3 show that drift increases *V_g_*, *L* and *δ* in regimes where *p_i_q_i_* tends to stay small at most loci at the deterministic equilibrium (*s̅* = 10_−2_, 10^−3^ in Figure 1), and has the opposite effect in regimes where *p_i_q_i_* is high (*s̅* = 10^−4^ in Figure 1). Simple approximations can be obtained when the effect of selection is negligible at most loci (see Supplementary File S2), which provide accurate predictions when *N_e_s̅* is sufficiently low, or when *s̅* ≪ *u* so that *p_i_* = 1/2 at most loci at the deterministic equilibrium (Figures S1 – S4). In this mutation-drift regime, *V_g_*, *L* and *δ* are nearly independent of *σ* when *N_e_u* ≪ 1 (the increase in variance caused by excess homozygosity being exactly compensated by the reduction in variance caused by the lower effective population size), or increase with *σ* for larger values of *N_e_u*. The discrepancies between analytical and simulation results observed in Figure 1 at high selfing rates are partly due to the reduction in effective population size *N_e_* caused by background selection, which is not accounted for in the diffusion model. An estimation of *N_e_* using the equilibrium diversity at a neutral locus (with an infinite number of possible alleles) at the mid-point of the chromosome (as in Roze, 2016) yielded an *N_e_* of approximately 740, 300 and 200 for *s̅* = 10^−4^, 10^−3^ and 10^−2^ (respectively) for *N* = 5,000 and *σ* = 1 (right-most points in Figure 1B). Replacing *N* by *N_e_* (1 + *F*) in the diffusion model provides predictions that closely match the simulation results for *s̅* = 10^−4^ and 10^−3^, suggesting that the initial discrepancy was indeed caused by background selection reducing *N_e_* (results not shown). However, for *s̅* = 10^−2^, the diffusion model still performs poorly despite the corrected *N_e_*. This implies that the discrepancy between analytical and simulation results is more likely due to interactions among loci, and possibly also to deviations of mean phenotypes from the optimum caused by genetic drift (that are not taken into account in the analysis).

**Figure 1.**
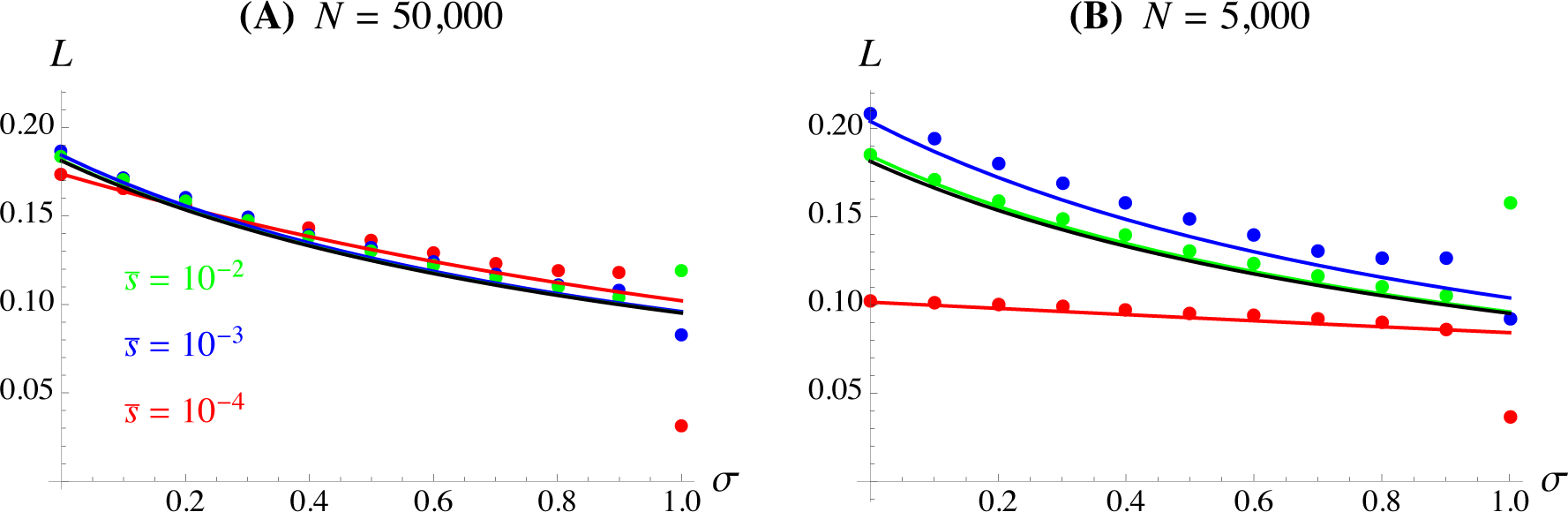
Mutation load *L* as a function of the selfing rate *σ*. Black curve: approximation for mutation-selection regime neglecting genetic associations (equation 24). The different colors correspond to different values of *s̅* as shown in A. Colored solid curves: results from the diffusion model (Supplementary File S2). Dots correspond to simulation results; in this and the following figures, error bars (computed by splitting the last 70,000 generations into 7 batches of 10,000 generations and calculating the standard error over batches) are smaller than the size of symbols in most cases. Other parameter values are *U* = 0.1, *R* = 20, *n* = 50, *m* = 5, *ℓ* = 10,000.

In Supplementary File S3, we derive expressions for the genetic variance, mutation load and inbreeding depression (for both the mutation-selection and the mutation-drift regimes) under the generalized fitness function given by equation 15. In the mutation-selection regime (*s̅_i_* ≫ 1/*N_e_*, *u* at most loci), one obtains:

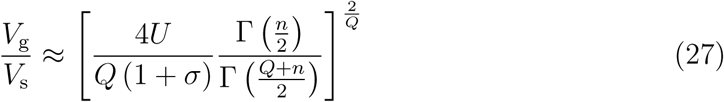

(where Γ is Euler’s Gamma function), while

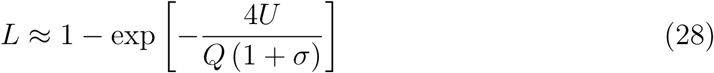

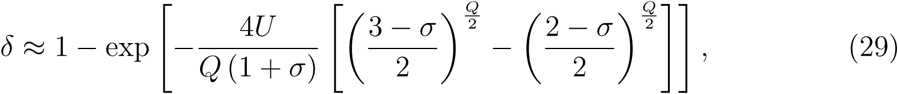

these equations being equivalent to equations 22, 24 and 26 when *Q* = 2. As shown by Figure 2, equations 27 – 29 provide good predictions of the simulation results when the population size and number of loci are sufficiently large (and selfing is not too high). As *Q* increases, the fitness peak becomes flatter around the optimum, and the equilibrium genetic variance increases (Figure 2B). However, despite increasing the genetic variance, higher values of *Q* lead to lower mutation loads due to the fact that deleterious alleles are more often eliminated when present in combination within the same genome: this corresponds to the classical result that negative epistasis reduces the mutation load in sexually reproducing populations (e.g., Kimura and Maruyama, 1966; Kondrashov and Crow, 1988). Indeed, the average epistasis between deleterious alleles equals zero for *Q* = 2, but becomes negative when *Q* > 2, and positive when *Q* < 2 (Gros et al., 2009). By contrast, inbreeding depression is less affected by *Q*, *δ* slightly increasing or decreasing as *Q* increases, depending on the selfing rate.

**Figure 2.**
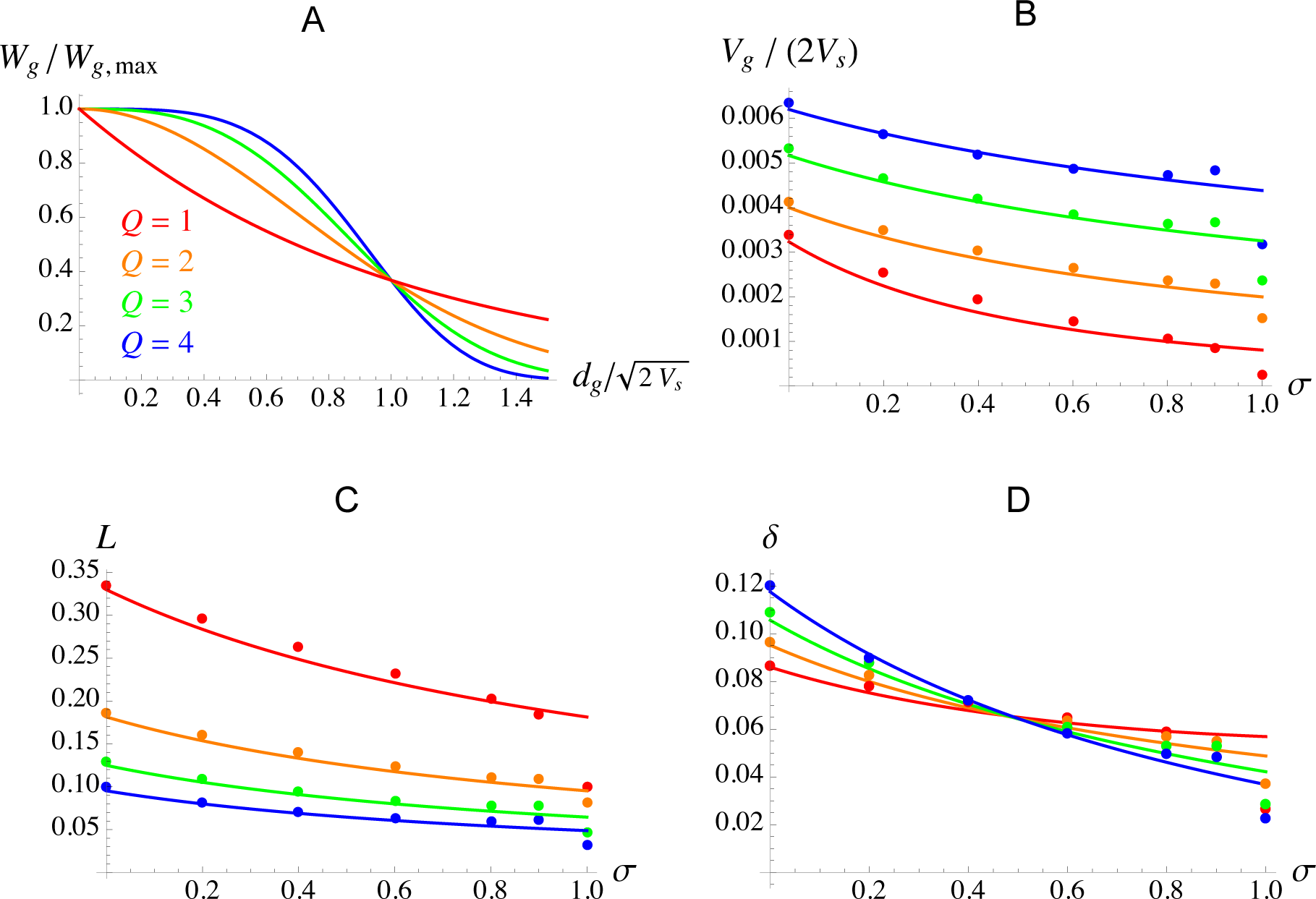
A: fitness as a function of the (scaled) distance from the optimum, for different values of the parameter *Q* (from equation 15). B, C, D: scaled genetic variance, mutation load and inbreeding depression as a function of the selfing rate *σ*, for different values of *Q*. The curves represent the analytical results (neglecting associations between loci) at mutation-selection balance (equations 27 – 29), while the dots correspond to simulation results. Parameter values: *N* = 50,000, *ℓ* = 10,000, *U* = 0.1, *R* = 20, *n* = 50, *m* = 5, *a*^2^/(2*V*_s_) = 0.0002 (yielding *s̅* = 0.001 for *Q* = 2).

Figure 3 shows the effects of the parameters *m* and *n* (for *Q* = 2). The degree of pleiotropy *m* of mutations affects their distribution of fitness effects (e.g., Lourenço et al., 2011). In Figure 3A, *s̅* is kept constant by decreasing the variance of mutational effects *a*^2^ as *m* increases (see equation 14). Increasing *m* (while keeping *s̅* constant) decreases the variance in fitness effects of mutations: indeed, one can show that the variance of mutational effects on log fitness (at the optimum) is given by 2*s̅*^2^/*m*. Figure 3A shows that *m* = 5 and *m* = 50 yield very close results when *s̅* = 10^−4^, as selection has a negligible effect at most loci for both values of *m* (for the parameter values used here), and the genetic variance does not depend on *m* at mutation-drift equilibrium (see equation B8 in Supplementary File S2). When *s̅* = 10^−2^, most loci are at mutation-selection balance (*s_i_* ≫ 1/*N*, *u*) for both *m* = 5 and *m* = 50, and the genetic variance is again not affected by *m* (see equation 22). Slightly different results are obtained for *m* = 1, due to the higher variance in fitness effects of mutations, causing a larger fraction of loci to be substantially affected by both selection and drift (this effect being captured by the diffusion model). Similarly, the effect of *m* is more pronounced when *s̅* = 10^−2^ and *σ* = 1, as *N_e_* is greatly reduced by background selection when selfing is high, causing higher proportions of loci to be substantially affected by drift.

**Figure 3.**
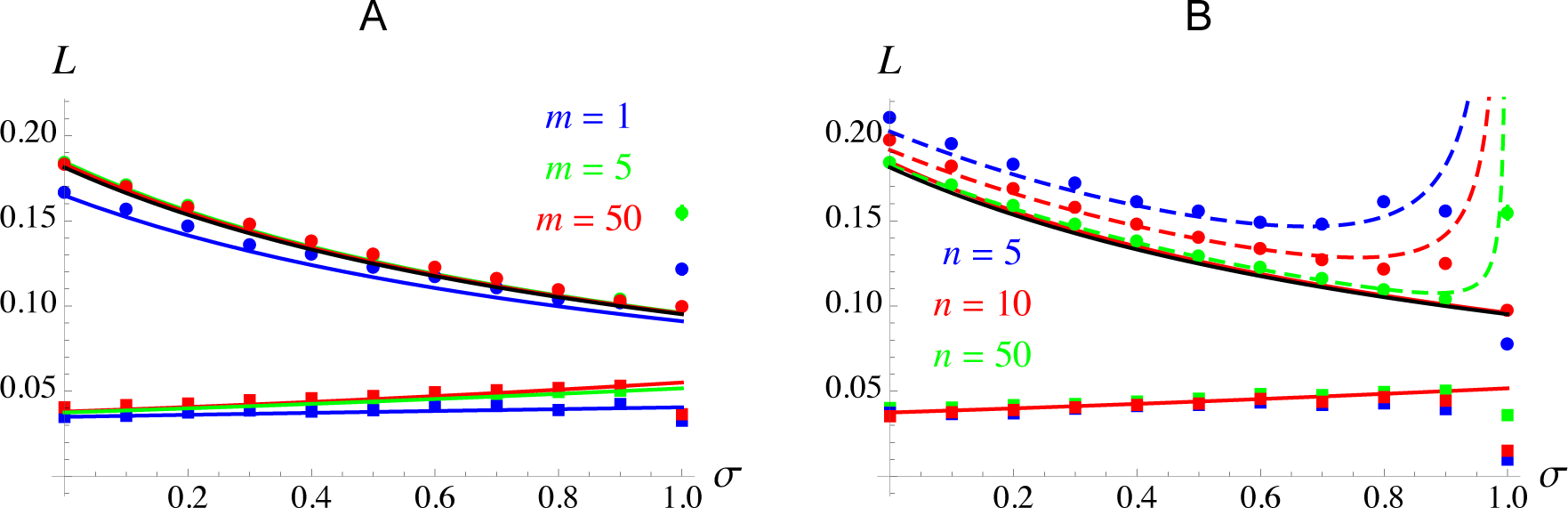
Mutation load *L* as a function of the selfing rate *σ* for *s̅* = 10^−2^ (top, filled circles) and *s̅* = 10^−4^ (bottom, filled squares). A: the different colors correspond to different values of *m* (degree of pleiotropy of mutations); B: the different colors correspond to different values of *n* (number of selected traits). Black curve: approximation for mutation-selection regime, neglecting genetic associations (equation 24). Colored solid curves: results from the diffusion model (Supplementary File S2). Colored dashed curves (in B): approximation including the effect of pairwise interactions among loci (equations 37, 39 and 40). Other parameter values are *U* = 0.1, *R* = 20, *N* = 5,000, *ℓ* = 1,000, *n* = 50 (in A), *m* = 5 (in B).

As shown by Figure 3B, the number of selected traits *n* has only little effect on the load in the mutation-drift regime (*s̅* = 10^−4^), in agreement with equation B9 in Supplementary File S2. However, while the diffusion model also predicts very little effect of *n* in the mutation-selection regime (*s̅* = 10^−2^), larger effects are observed in the simulations, with larger deviations from the analytical predictions (and higher load) for lower values of *n*. These deviations are caused by associations between loci (which are neglected in equation 24 and in the diffusion model). In the next subsection, we show that the relative effect of these associations is indeed stronger when *n* is lower, and derive an approximation including the effect of pairwise genetic associations that better matches the simulation results.

### Effects of associations between loci

In Supplementary File S1, we derive approximations for the effects of associations between pairs of loci on the genetic variance at mutation-selection balance, under a Gaussian fitness function (*Q* = 2). For this, we assume that these associations remain weak, and neglect the effects of all associations involving more than two loci. As shown by equation 11, associations *D*̃*_ij_*, *D*̃*_i,j_* (between alleles at loci *i* and *j*, either on the same or on different haplotypes) directly affect the genetic variance. At equilibrium, these associations are approximately given by:

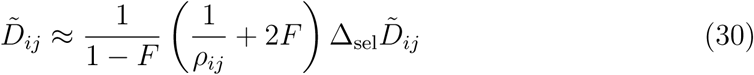

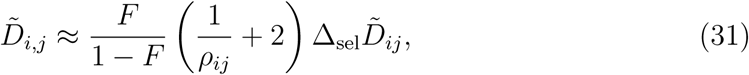

where again *F* = σ/(2 − *σ*), *ρ_ij_* is the recombination rate between loci *i* and *j*, and Δ_sel_*D*̃*_ij_* is the change in *D*̃*_ij_* and *D*̃*_i,j_* due to selection:

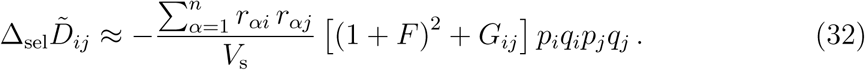

The term *G_ij_* in equation 32 represents the identity disequilibrium between loci *i* and *j*: the correlation in identity by descent between loci, generating a correlation in homozygosity (Weir and Cockerham, 1973, Supplementary File S1). Equation 32 shows that stabilizing selection generates a positive association between alleles at loci *i* and *j* that tends to displace phenotypes in opposite directions (allele 1 with allele 1, and allele 0 with allele 0 if 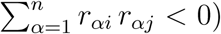): the effect of the deleterious allele at locus *i* is then partially compensated by its associated allele at locus *j* (e.g., Bulmer, 1974; Lande, 1976; Turelli and Barton, 1990). This effect of selection is strengthened by homozygosity (and correlations in homozygosity between loci) caused by selfing. As may be seen from equations 30 and 31, *D*̃*_i,j_ ≈ F D*̃*_ij_* when loci are tightly linked (*ρ_ij_ ≪* 1), as expected from separation of timescales arguments (e.g., Nordborg, 1997; Roze, 2016). However, our approximations diverge as recombination tends to zero (or as the selfing rate tends to 1), due to the assumption that genetic associations remain weak.

We show in Supplementary File S1 how *D*̃*_ij_* and *D*̃*_i,j_* can be summed over all pairs of loci in order to compute their overall direct effect on the genetic variance (second term of equation 11). These associations depend on recombination rates through the terms in 1*/ρ_ij_* in equations 30 and 31, and also through the identity disequilibrium *G_ij_* in equation 32. However, because *G_ij_* only weakly depends on the recombination rate, its average over all pairs of loci is often very close to the value obtained under free recombination, provided that the genome map length is not too small (see Supplementary Figure S5). In the following, we thus approximate *G_ij_* by its expression for freely recombining loci, denoted *G*:

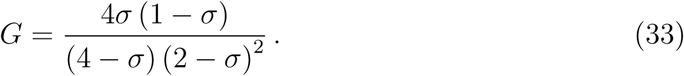

By contrast, linkage has more effect on the average of 1/*ρ_ij_* over all pairs of loci, corresponding to the inverse of the harmonic mean recombination rate between all pairs of loci (denoted *ρ*_H_ thereafter). Assuming that the number of loci is large, one obtains for the direct effect of linkage disequilibria on the genetic variance (see Supplementary File S1):

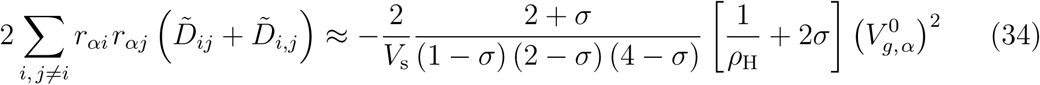

where the genic variance 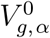 may be replaced by its expression to leading order, given by equation 21. Equation 34 shows that the immediate effect of associations between alleles with compensatory phenotypic effects is to reduce the genetic variance (since this term is negative). The fraction in equation 34 is an increasing function of *σ*, which implies that self-fertilization increases the strength of associations, thus decreasing *V*_g*, α*_. However, because the genic variance is expected to decrease with *σ* (equation 21), the direct effect of linkage disequilibria on *V*_g*, α*_ may remain approximately constant (or even slightly decrease) as *σ* increases from zero.

Associations between loci do not only affect *V*_g*, α*_ through equation 34, however, but also affect the equilibrium allele frequencies and the excess homozygosity *D_i,i_* at each locus. The effect on *D_i,i_* is mainly driven by identity disequilibria: indeed, neglecting associations between 3 or more loci, one obtains (see Supplementary File S1):

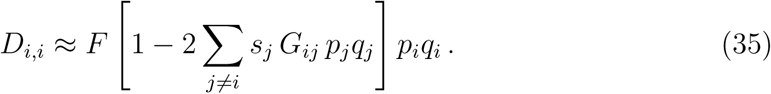

Equation 35 is equivalent to equation 5 in Roze (2015) (which is expressed to the first order in *p_j_*), as can be noted by replacing *s* and *h* in Roze (2015) by 4*s_j_* and 1/4. It shows that identity disequilibria reduce the excess homozygosity at each locus: this is due to the fact that homozygotes at locus *i* are more likely to be also homozygous at locus *j*, while homozygotes at locus *j* have a lower fitness than heterozygotes when deleterious alleles are partially recessive. Identity disequilibria thus tend to reduce the genetic variance through this effect on *D_i,i_*, by an amount corresponding to the sum of the term in *G_ij_* in equation 35 over all pairs of loci. Approximating *G_ij_* by its expression for freely recombining loci (equation 33), one obtains that this effect reduces *V* by approximately 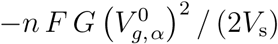, where again 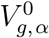 may be replaced by the expression given by equation 21, to leading order (see Supplementary File S1).

Finally, associations between loci affect equilibrium allele frequencies (*p_i_q_i_*) at each locus. As shown in Supplementary File S1, both the linkage disequilibria generated by epistasis and the identity disequilibria caused by partial selfing reduce the efficiency of purging, thereby increasing *p_i_q_i_* and thus the genic variance. Indeed, an expression for the effect of selection on *p_i_q_i_* that includes the effects of pairwise associations is, to leading order:

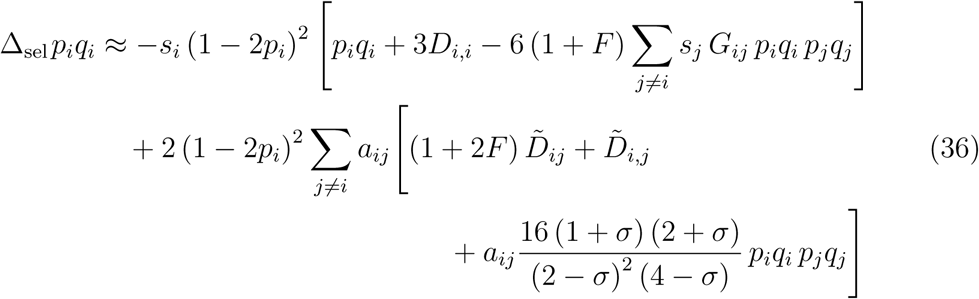

with *a_ij_* = −Σ*_α_r_αi_r_αj_*/(2*V_s_*), and where *D*̃*_ij_*, *D*̃*_i,j_* and *D_i,i_* are given by equations 30, 31 and 35. The first line of equation 36 is equivalent to equation 6 in Roze (2015), showing that identity disequilibria reduce the efficiency of purging by decreasing the excess homozygosity (*D_i,i_*), and by two additional effects represented by the term in 6 (1 + *F*) (see Roze (2015) for interpretation of these effects). The term on the second and third lines (proportional to 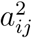) represents the effect of epistasis between loci: this term also reduces purging, since selection against deleterious alleles is less efficient when these alleles are partially compensated by alleles at other loci.

An expression for the genic variance at mutation-selection balance is given by equation A65 in Supplementary File S1. From this, one obtains for the genetic variance:

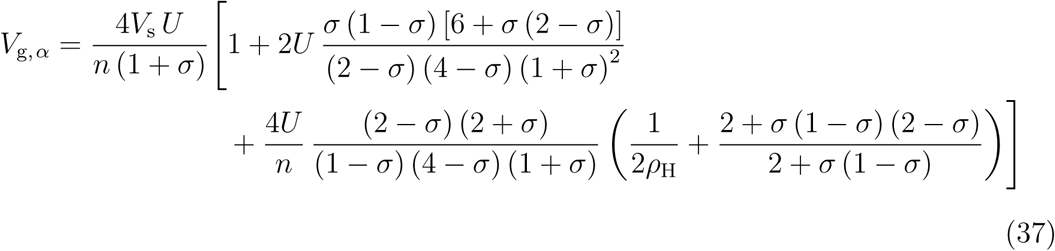

where the terms in *U* between the brackets correspond to the effect of between-locus associations. The first of these terms (on the first line of equation 37) represents the effect of identity disequilibria, while the term in *U*/*n* on the second line represents the effect of epistasis (compensatory effects between alleles at different loci). Both terms are positive, indicating that the overall effect of interactions between loci is to increase the genetic variance, due to the fact that correlations in homozygosity and compensatory effects between mutations both reduce the efficiency of purging (equation 36). Furthermore, while the effect of identity disequilibria scales with *U*, the effect of epistasis scales with *U*/*n*: indeed, it becomes less and less likely that alleles at different loci have compensatory effects on all of the traits as the dimensionality of the fitness landscape increases. Finally, the effect of epistasis is more strongly affected by linkage between loci (through the term in 1*/ρ*_H_); the effect of linkage on the term in *U* representing the effect of linkage disequilibria is weaker, and has been neglected in equation 37. Under random mating (*σ* = 0), equation 37 simplifies to:

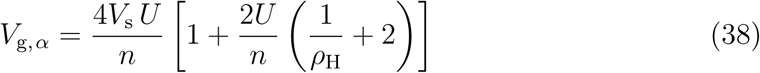

which takes a similar form as equation 4.16 in Turelli and Barton (1990) in the case of a single selected trait (*n* = 1). Supplementary Figure S6 shows how the equilibrium genetic variance and its different components vary with the selfing rate, in a regime where both identity disequilibria and epistasis have significant effects (*U* = 1).

Assuming that 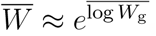, an approximation for the load at mutation-selection balance is 1 − exp[−*n V_g_*/(2*V_s_*)], where *V_g_* is the genetic variance given by equation 37 (the same for all traits). A slightly better approximation can be obtained by using 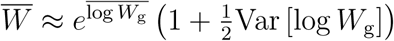, where Var [log*W_g_*] is the variance in log fitness in the population (Roze, 2015). To leading order, it is given by (see Supplementary File S1):

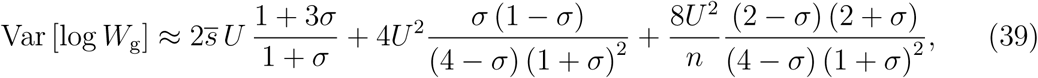

simplifying to 2*U* (*s̅* + 4*U*/*n*) in the absence of selfing. The first term of equation 39 represents the sum of single-locus contributions to the variance in log fitness, while the second and third term correspond to the effects of identity disequilibria and epistasis (respectively), both increasing the variance in fitness. The mutation load is then given by:

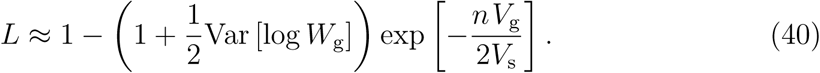

Similarly, we show in Supplementary File S1 that an expression for inbreeding depression including the effects of pairwise associations between loci is:

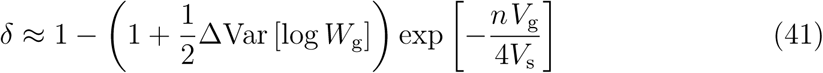

where ΔVar [log *W_g_*] is the difference in variance in log fitness between selfed and outcrossed offspring, given by:

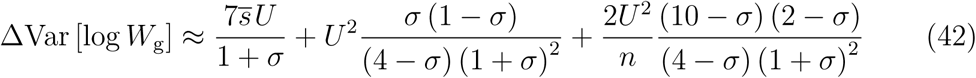

(the terms in Var [log *W_g_*] and ΔVar [log *W_g_*] in equations 40 and 41 are often small, however, and may thus be neglected). After replacing *V_g_*, Var [log *W_g_*] and ΔVar [log *W_g_*] by the expressions given by equations 37, 39 and 42, the approximations obtained for the load and inbreeding depression include terms in *U*^2^ representing the effect of identity disequilibria, and terms in *U*^2^/*n* representing the effect of epistasis between loci. The terms in *U*^2^ are identical to the terms representing the effect of identity disequilibria in a model with purely multiplicative selection against deleterious alleles (no epistasis) when setting the dominance coefficient *h* of deleterious alleles to 1/4 (equations 11 and 14 in Roze, 2015). The novelty here thus corresponds to the effect of epistasis (compensatory effects between deleterious alleles), that tends to increase *V_g_*, *L*, *δ* by reducing the efficiency of purging.

Figure 3B shows that equations 37, 39 and 40 capture the increase in load observed in the simulations as the number of traits *n* decreases (see Supplementary Figure S7 for the genetic variance and inbreeding depression). Note that the harmonic mean recombination rate *ρ*_H_ between pairs of loci under our simulated genetic architecture (linear chromosome with equally spaced loci) can be obtained from:

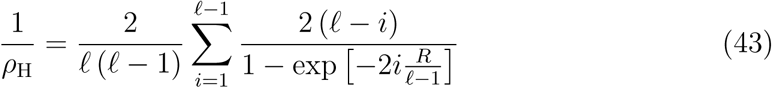

(see Appendix 2 in Roze and Blanckaert, 2014), yielding *ρ*_H_ *≈* 0.42 for *ℓ* = 1,000 and *R* = 20. Figure 4 shows that for low or moderate selfing rates, decreasing the genome map length from *R* = 20 to *R* = 1 increases the mutation load, by increasing the strength of linkage disequilibria caused by epistasis, that in turn reduce the efficiency of purging. In this regime, equations 37, 39 and 40 provide an accurate prediction for the load (see Supplementary Figure S8 for genetic variance and inbreeding depression). At high selfing rates, however, a different regime is entered, in which the assumption of weak genetic associations breaks down. As can be seen in Figure 4, in this regime (which spans a broader parameter range under tighter linkage) the load decreases more rapidly as *σ* increases. Increasing linkage tends to reduce the mutation load when the selfing rate is high, although the effect of *R* vanishes when *σ* = 1. When linkage is extremely tight, the approximations given above break down for all values of *σ*: as shown by Figures 4 and S8, decreasing *R* has a non-monotonic effect on the genetic variance, load and inbreeding depression when selfing is small to moderate, the lowest values of *V_g_*, *L* and *δ* being reached when *R* = 0 (in which case selfing has no effect). An approximation for the genetic variance under complete linkage can be obtained by treating the whole genome as a single locus with a very large number of possible alleles, and assuming a Gaussian distribution of allelic effects in the population (Lande, 1977; Supplementary File S4). This yields:

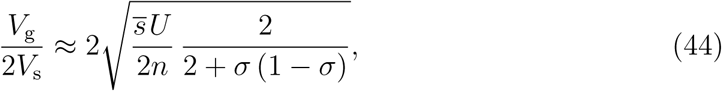

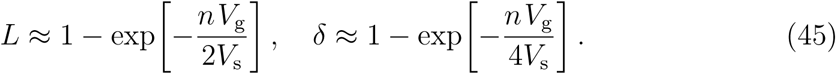

Note that equation 44 is equivalent to equation 3A in Charlesworth and Charlesworth (1995) when *σ* = 1 and *n* = 1. As shown by Figures 4 and S8, equations 44 and 45 only slightly overestimate *V_g_*, *L* and *δ* when *σ* = 1 and/or *R* = 0. As shown below, better predictions are observed for higher values of *U*/*n* and lower values of *s̅*.

**Figure 4.**
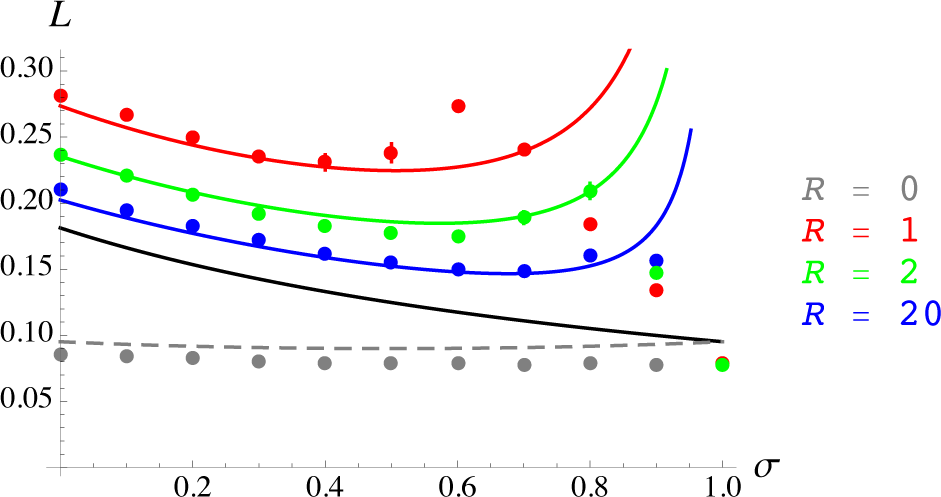
Mutation load *L* as a function of the selfing rate *σ*, for *s̅* = 0.01, *n* = *m* = 5 and different values of the genome map length *R*, yielding (using equation 43) *ρ*_H_ ≈ 0.07, 0.13 and 0.42 for *R* = 1, 2, 20 (respectively). Dots: simulation results; black curve: approximation for mutation-selection regime neglecting genetic associations (equation 24); colored curves: approximation including the effect of pairwise interactions among loci (equations 37, 39 and 40); dashed grey curve: single-locus model with many alleles, assuming a Gaussian distribution of allelic values (equations 44 and 45). Other parameter values are as in Figure 3.

The effects of identity disequilibria between loci remain negligible for the parameter values used in Figures 3 and 4. As shown by Figure 5, identity disequilibria become more important for higher values of the mutation rate *U*. Indeed, the relative effects of identity disequilibria on the load can be deduced from the differences between the three curves in each panel of Figure 5, the red curves showing the predicted mutation load in the absence of epistasis, but taking into account identity disequilibria (obtained by removing the terms in *U*^2^/*n* from equations 37, 39 and 40, leading to an expression equivalent to equation 11 in Roze, 2015). The difference between the black and red curves thus represents the predicted effect of identity disequilibria on the load, while the difference between the red and green curves corresponds to the additional effect of epistasis. Simulations indicate that the change in regime observed above a threshold selfing rate (around *σ* = 0.5 for *U* = 1 in Figure 5) is due to epistasis, since this threshold is not observed in simulations without epistasis (red dots). Supplementary Figure S9 shows that this threshold pattern is little affected by population size *N*, as long as the effects of drift remain small. Similarly, the results only weakly depend on the number of loci *ℓ*, as long as the mutation rate per locus *u = U/ℓ* is small enough so that *p_i_q_i_* < 1/4 at most loci (see Supplementary Figure S10 for distributions of allele frequencies in simulations with *ℓ* = 1,000 and *ℓ* = 10,000). Figure S9 also shows that the results are little affected by the degree of pleiotropy of mutations *m*, as long as *s̅* remains constant. However, *s̅* does affect *V_g_*, *L* and *δ* in the regime where our approximations break down. As shown by Figure 6, decreasing *s̅* lowers the threshold selfing rate above which our approximations are not valid and results in lower equilibrium mutation loads (see Supplementary Figure S11 for results on *V_g_* and *δ*). Figures 6 and S11 also show that, when *s̅* is sufficiently small, the single-locus, Gaussian model (equations 44 and 45, dotted curves on the figures) provides accurate predictions for *V_g_*, *L* and *δ* under complete selfing (*σ* = 1).

**Figure 5.**
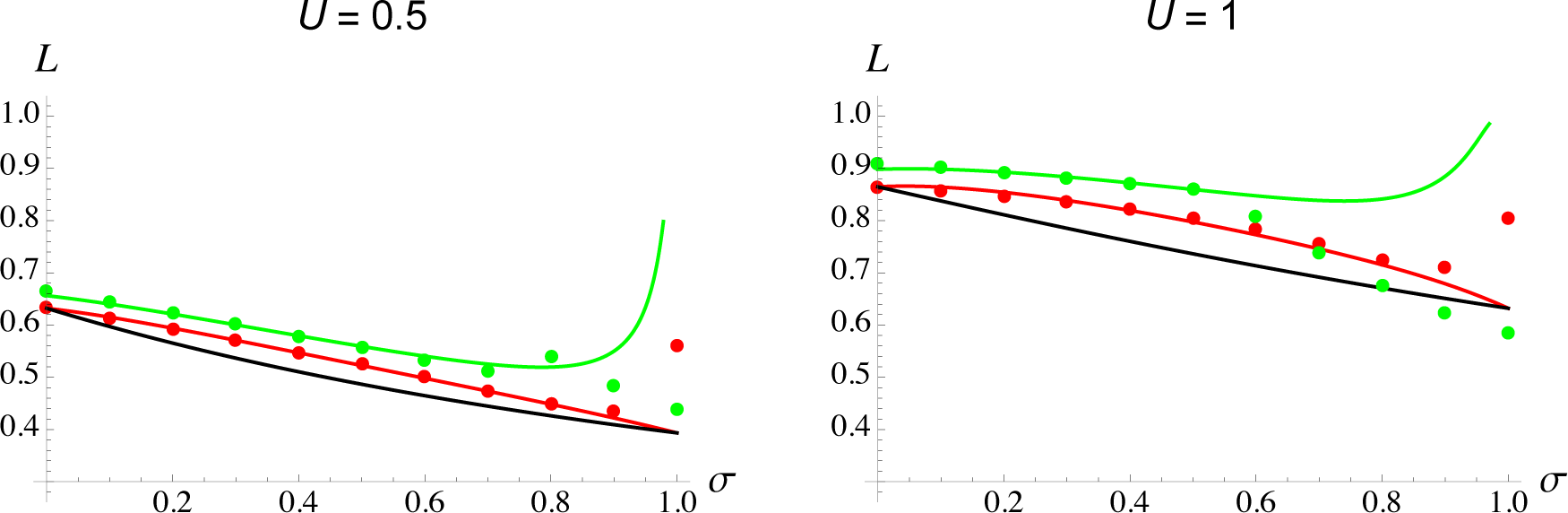
Mutation load *L* as a function of the selfing rate *σ*, for *s̅* = 0.01, *n* = 50, *m* = 5, *U* = 0.5 (left) and 1 (right). The black curves correspond to the approximation for mutation-selection regime, neglecting genetic associations (equation 24). Green curves: approximation including the effect of pairwise interactions among loci (equations 37, 39 and 40); red curves: approximation including the effects of identity disequilibria between loci, but not the effects of epistasis (obtained by removing the terms in *U*^2^/*n* from equations 37, 39 and 40, equivalent to equation 11 in Roze, 2015). Green dots: simulation results; red dots: results from the simulation program used in Roze (2015) representing multiplicative selection (no epistasis), with *s* = 0.04 and *h* = 0.25. Other parameter values are *N* = 5,000, *ℓ* = 10,000, *R* = 20 (yielding *ρ_H_* ≈ 0.38).

**Figure 6.**
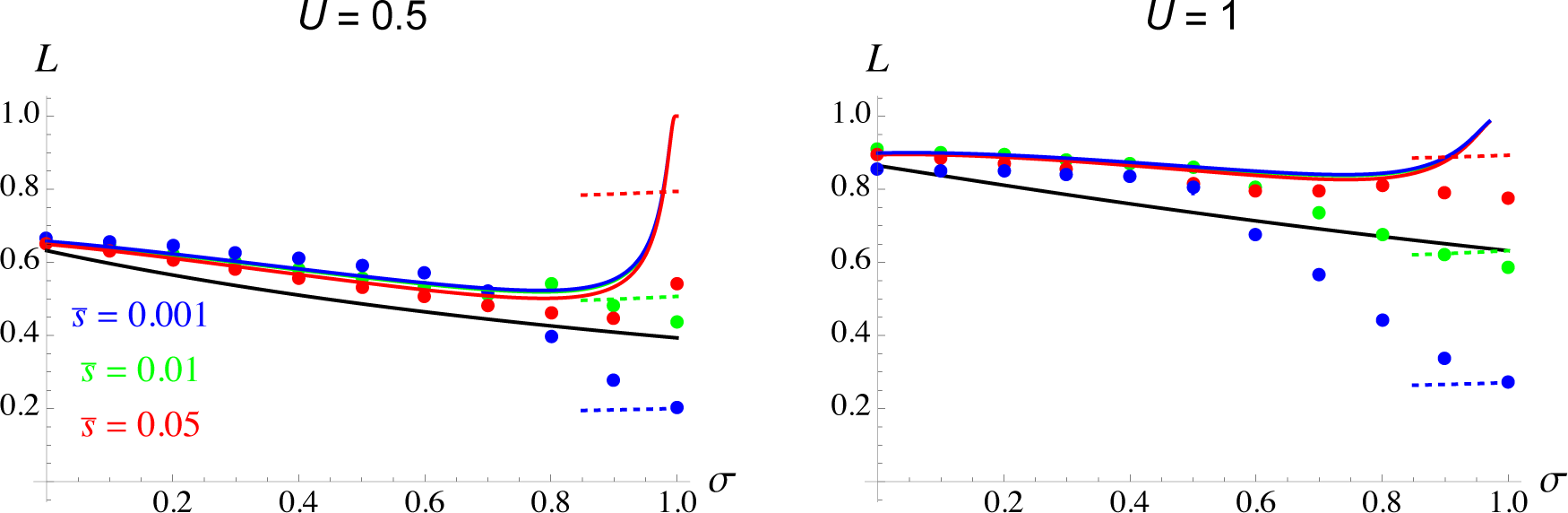
Mutation load *L* as a function of the selfing rate *σ*, for different values of the mutation rate *U* and average heterozygous effect of mutations *s̅*; other parameter values are as in Figure 5. Dots: simulation results; black curves: approximation for mutation-selection regime, neglecting genetic associations (equation 24); solid colored curves: approximation including the effect of pairwise interactions among loci (equations 37, 39 and 40); dotted colored curves: single-locus model with many alleles, assuming a Gaussian distribution of allelic values (equations 44 and 45).

In Figure 7 we show that decreasing the number of traits under selection *n* decreases the threshold selfing rate above which our approximations break down (see Supplementary Figure S12 for inbreeding depression and scaled genetic variance). Below the threshold, the mutation load decreases as *n* increases, as predicted by our analytical results (although our approximations become less precise for low *n* and high *U*), while *n* has the opposite effect above the threshold. Overall, we observe that in this second regime (in which interactions between loci have important effects), the mutation load generally increases with the number of selected traits, the fitness effects of mutations *s̅*, the mutation rate *U* and recombination rate (through the parameter *R*). However, Figure 8 shows that the effects of these parameters on inbreeding depression are more complicated. In particular, outbreeding depression (negative *δ*) may occur in regimes where the effects of epistasis are particularly strong (high *U*, low *n*) and when the selfing rate is moderate to high (above 0.5 but below 1), outbreeding depression becoming stronger when *s̅*, *U* and *R* increase (the approximation derived from equation 37 fails for all values of *σ* for the parameter values used in Figure 8, and is not shown here). Supplementary Figures S13 and S14 show that for the same parameter values, *V_g_* and *L* always increase when *s̅*, *U* and *R* increase.

**Figure 7.**
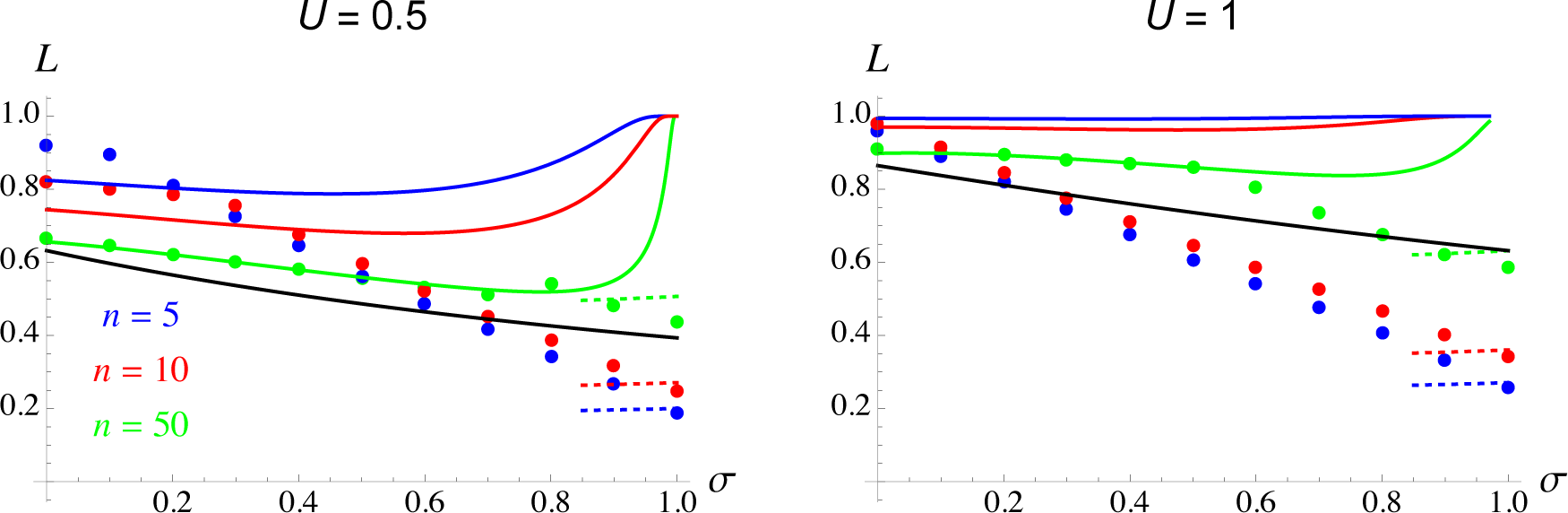
Mutation load *L* as a function of the selfing rate *σ*, for different values of the mutation rate *U* and number of selected traits *n*; other parameter values are as in Figure 5. Dots: simulation results; black curves: approximation for mutation-selection regime, neglecting genetic associations (equation 24); solid colored curves: approximation including the effect of pairwise interactions among loci (equations 37, 39 and 40); dotted colored curves: single-locus model with many alleles, assuming a Gaussian distribution of allelic values (equations 44 and 45).

**Figure 8.**
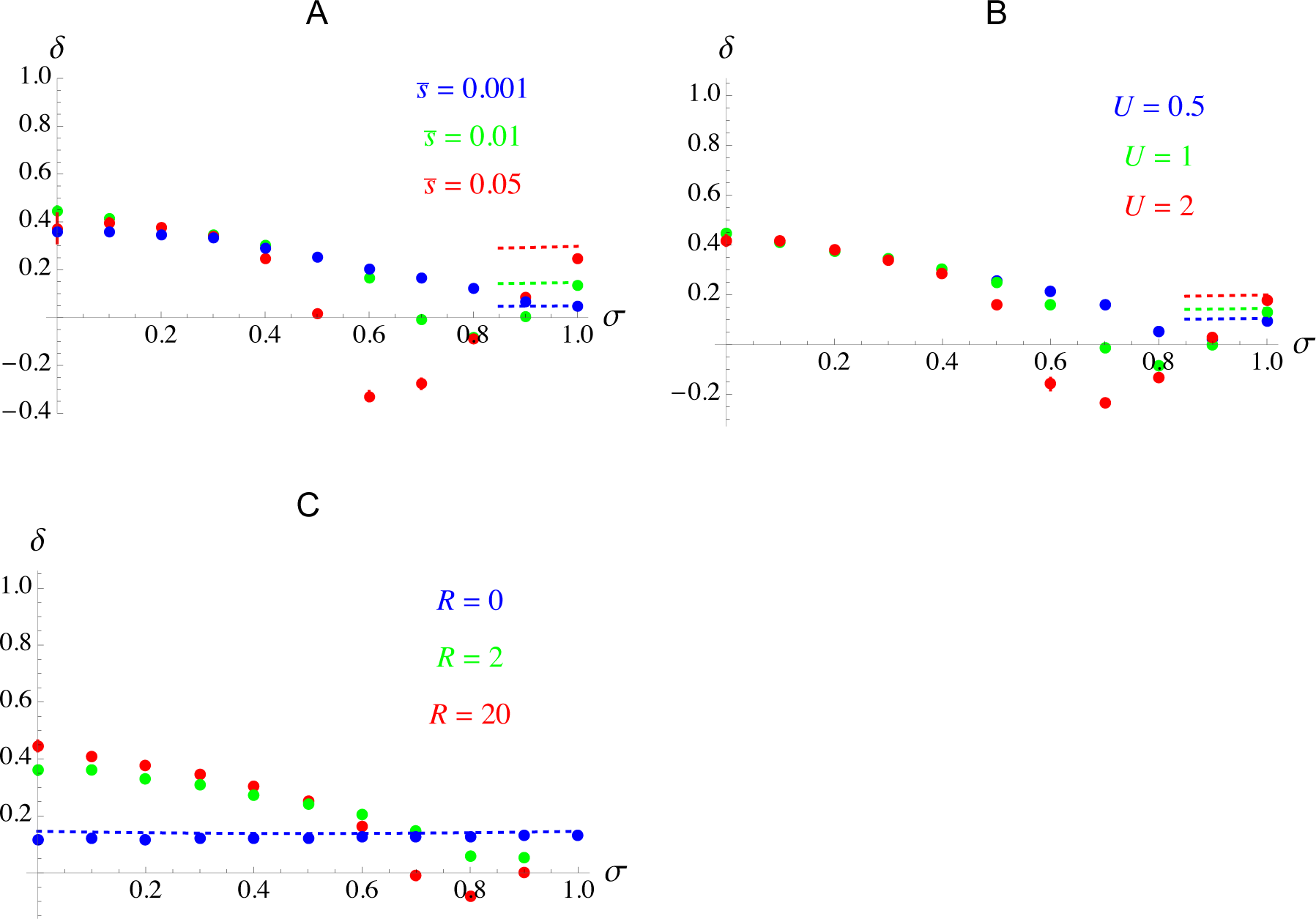
Inbreeding depression *δ* as a function of the selfing rate *σ*, for *n* = *m* = 5 and different values of *s̅* (A), *U* (B) and *R* (C). Dots: simulation results; dotted curves: single-locus model with many alleles, assuming a Gaussian distribution of allelic values (equations 44 and 45). Other parameter values are *N* = 5,000, *ℓ* = 10,000, *R* = 20, *U* = 1 and *s̅* = 0.01.

## DISCUSSION

The response of a population to environmental change depends critically on its genetic diversity. Our results predict that the level of genetic variation maintained at equilibrium under stabilizing selection acting on quantitative traits is generally lower in more highly selfing population, due to more efficient purging (although increasing selfing may sometimes increase genetic variation, for example when mutations have weak fitness effects, as shown by Figure S3). This finding agrees with Charlesworth and Charlesworth’s (1995) theoretical prediction that fully selfing populations should maintain lower genetic variance for quantitative traits under stabilizing selection than fully outcrossing ones, and with several empirical studies comparing levels of genetic variation for morphological traits in closely related pairs of plant species with contrasted mating systems (Charlesworth and Charlesworth, 1995; Geber and Griffen, 2003; Bartkowska and Johnston, 2009 and references therein). We also show that the lower level of variation present in more highly selfing populations is associated with lower values of the mutation load and inbreeding depression. The meta-analysis carried out by Winn et al. (2011) showed that inbreeding depression is indeed lower in highly selfing plant species compared to species with lower selfing rates, while no significant difference is observed between species with low vs. intermediate selfing rates. It has been put forth that correlations in homozygosity between selected loci may suppress purging at moderate selfing rates (“selective interference”, Lande et al., 1994; Winn et al., 2011); this, however, would imply that a large number of segregating deleterious alleles have very low dominance coefficients, generating very high inbreeding depression (Kelly, 2007; Roze, 2015), which seems unlikely. Another possible explanation for the lack of purging at intermediate selfing rates involves epistasis (compensatory effects between mutations coding for the same quantitative trait, Lande and Porcher, 2015). Our analysis of the effects of epistasis (under assumptions that differ from those made in Lande and Porcher’s model) shows that different regimes are possible, and outlines how the parameters affect transitions between these regimes.

In our model, the effect of epistasis on the equilibrium genetic variance *V_g_* is inversely proportional to effective recombination rates between selected loci, and scales with *U*/*n* (where *n* is the number of selected traits and *U* the total mutation rate on those traits). Indeed, *U*/*n* determines the number of segregating “interacting” mutations, that is, mutations with epistatic fitness effects. As *n* tends to infinity, all mutations become orthogonal in phenotypic space (with independent fitness effects), and our results converge to the results from previous population genetics models without epistasis (e.g., Charlesworth and Charlesworth, 1987; Roze, 2015). When *U*/*n* is small and map length *R* is sufficiently large, associations between loci have little effect. Under Gaussian stabilizing selection (*Q* = 2), the average coefficient of epistasis between mutations (on fitness) is zero (Martin and Lenormand, 2006b) while the dominance coefficient of deleterious alleles (in an optimal genotype) is close to 0.25 under the assumption of additive effects on phenotypes. In this case, we found that classical deterministic expressions based on single-locus models (hence neglecting the variance in epistatic interactions) provide accurate predictions for the mutation load *L* and inbreeding depression *δ*. Simple approximations are also obtained under the more general fitness function given by equation 15, confirming that the mutation load is an increasing function of the average coefficient of epistasis between mutations (Kimura and Maruyama, 1966; Kondrashov and Crow, 1988; Phillips et al., 2000; Roze and Blanckaert, 2014). Neglecting the effect of associations between loci also allowed us to explore the effects of drift using diffusion methods. As in previous studies (e.g., Charlesworth, 2013; Roze and Blanckaert, 2014), we found that drift may lower the mutation load by reducing *V_g_*. However, this result probably strongly depends on the assumption that mutations may increase or decrease phenotypic traits with the same probability (no mutational bias): indeed, previous works showed that drift may increase the load in the presence of a mutational bias by displacing mean phenotypes away from the optimum (Zhang and Hill, 2008; Charlesworth, 2013). Given that partial selfing reduces effective population size, it would be of interest to study the combined effects of drift and mutational bias in models with selfing.

The variance in epistasis has stronger effects as the *U*/*n* ratio increases and/or as the effective recombination rate decreases (*i.e.* due to selfing). Our results showed that two different regimes are possible. (1) When genetic associations (linkage disequilibria) generated by epistasis stay moderate, the overall effect of epistasis is to increase *V_g_*, *L* and *δ* by decreasing the efficiency of selection against deleterious alleles. This regime is generally well described by our model taking into account the effects of associations between pairs of loci. This result bears some similarity with the result obtained by Phillips et al. (2000), showing that the variance in epistasis between deleterious alleles increases the mutation load. Equation 2.1 in Phillips et al. (2000) is not fully equivalent to our expression for the load under random mating, however, possibly due to different assumptions on the relative orders of magnitude of *s_i_*, *s_j_* and *e_ij_* (where *s_i_* and *s_j_* are the strength of selection at loci *i* and *j* and *e_ij_* is epistasis between those loci) and how they covary. Nevertheless, it is interesting to note that both results become equivalent if Var [*e_ij_ /* (*s_i_sj*)] in Phillips et al. (2000) is replaced by Var [*eij*/*s̅*^2^] (where Var stands for the variance across all pairs of loci), using the fact that Var [*eij*] = 4*s̅*^2^/*n* in our model (with a Gaussian fitness function). (2) Increasing the value of *U*/*n* and/or reducing effective recombination rates or *s̅* generates a transition to a different regime in which the effect of the variance in epistasis switches, reducing *V*g, *L* and *δ*. Because our analytical approach fails in this regime (presumably due to higher-order associations between loci), it is more difficult to obtain an intuitive understanding of the selective mechanisms involved. However, it is likely that selection operates on multilocus genotypes (comprising combinations of alleles with compensatory effects) that can be maintained over many generations due to high selfing rates and/or low recombination. A similar transition from genic to genotypic selection as recombination decreases was described by Neher and Shraiman (2009), using a haploid model in which epistasis is randomly assigned to genotypes.

Although our results show some qualitative similarities with those obtained by Lande and Porcher (2015) — e.g., the same transition between regimes occurs in both models as selfing increases — several differences can be observed. In particular, Lande and Porcher’s model predict little or no effect of selfing on *V*g below the threshold selfing rate corresponding to the change in regime, and an abrupt change in *V*g at the threshold (except in their infinitesimal model). A step change such as this is never observed in our model, even for parameter values at which the effect of drift should be negligible at most loci. These differences between the models are not due to the different genetic architectures considered (biallelic vs. multiallelic): indeed, Supplementary Figures S15 and S16 show that assuming biallelic loci or an infinite number of possible alleles per locus in our individual-based simulations yields very similar results (for *ℓ* = 1,000 and *ℓ* = 10,000). Rather, they must be due to Lande and Porcher’s assumption of a Gaussian distribution of allelic effects maintained at each locus in each selfing age class, implicitly assuming a sufficiently high mutation rate per locus *u* and low fitness effect of mutations *s̅* (Turelli, 1984). In our multiallelic simulations (with *u* = 10^−5^ to 10^−3^ and *s̅* = 0.01), the number of alleles maintained at each locus is not sufficiently large to generate a Gaussian distribution of segregating allelic effects (see Figures S15 and S16). One may also note that the effect of the number of selected traits *n* seems different in both models (compare Lande and Porcher’s Figure 5 and 6 to our Figure 7), but this is due to the fact that the overall mutation rate *U* is proportional to *n* in Lande and Porcher’s model (while *U* is fixed in Figure 7). Increasing both *n* and *U* in order to maintain a constant *U*/*n* ratio, we indeed observed that the transition between regimes occurs at lower selfing rates when *n* is larger, as in Lande and Porcher’s Figure 5 and 6 (results not shown). In general, whether *U* should scale with *n* depends on the degree of pleiotropy of mutations (Lande and Porcher assume no pleiotropy). Our model allowed us to explore the effects of pleiotropy through the parameter *m*, showing that pleiotropy mostly affects the results through its effect on *s̅* (equation 14). The equilibrium genetic variance thus depends on *m* in regimes where *V*g is affected by *s̅*, in particular when *N_e_s̅* ≈ 1 or lower (Figures 1, S1 – S4), and when genetic associations are strong (Figure 6). However, pleiotropy may have stronger effects under different assumptions regarding the genetic architecture of traits, for example when different sets of traits are affected by different sets of loci (modular pleiotropy, Welch and Waxman, 2003). The effects of selective or mutational covariance among traits would also be interesting to explore: indeed, such covariances decrease the effective number of selected traits (Martin and Lenormand, 2006b), potentially increasing the importance of associations between loci.

In the regime where genetic associations generated by epistasis reduce *V*g (regime (2) mentioned above), outbreeding depression may occur due to the lower fitness of recombinants between selfing lineages maintaining coadapted gene complexes (Figure 8), a result shared with Lande and Porcher’s (2015) Gaussian Allele Model. In our additive model of phenotypic effects, outbreeding depression should only be expressed in F2 individuals (that is, among the offspring of an individual produced by a cross between different selfing lineages), once recombination has disrupted compensatory associations between alleles at different loci. This explains why outbreeding depression is not observed under complete (or nearly complete) selfing in Figure 8, as all outcrossed individuals are F1 hybrids between selfing lineages. Outbreeding depression between lineages collected from the same geographical location has been observed in highly selfing plants (Parker, 1992; Volis et al., 2011) and *Caenorhabditis* nematodes (Dolgin et al., 2007; Gimond et al., 2013). In all cases, estimated selfing rates are higher than those leading to *δ* < 0 in our simulations, however, and outbreeding depression was observed in F1 offspring of crosses between inbred lines of nematodes. The occurrence of outbreeding depression at higher selfing rates may be partly explained by the fact that experimental crosses were often performed between genetically different lines; by contrast, in our simulations the parents of an outcrossed individual may share the same genotype (in particular when the number of genetically different selfing lineages is reduced due to the low effective size of highly selfing populations), reducing the magnitude of outbreeding depression. However, the occurrence of outbreeding depression in F1 individuals must involve dominance effects which are absent from our model. Exploring the effects of dominance/recessivity of mutations on phenotypic traits would be an interesting extension of this work.

Due to the lower genetic diversity of self-fertilizing populations, it has been suggested that they should be less able to adapt to a changing environment (e.g., Stebbins, 1957; Williams, 1992; Takebayashi and Morrell, 2001). In the absence of epistasis, existing models indeed predict that selfing populations should have lower rates of adaptation than outcrossing ones (Gl´emin and Ronfort, 2013; Hartfield and Gl´emin, 2016). When compensatory effects between mutations are possible, however, a substantial amount of genetic variance may be hidden by genetic associations between loci in highly selfing populations (Lande and Porcher, 2015, the present study). After a change in environment, this variance may be liberated by rare outcrossing events, increasing the short-term evolutionary response of highly (but not fully) selfing populations. Exploring how selfing affects adaptation under directional selection, and more generally how the variability of epistatic interactions between loci may influence the evolution of mating systems represents a natural next step of this work.

## Acknowledgements

We thank Aneil Agrawal, Patrice David, Sylvain Gl´emin, Fr´ed´eric Guillaume, Thomas Lenormand, Emmanuelle Porcher, Oph´elie Ronce, Jo¨elle Ronfort and an anonymous reviewer for helpful discussions and comments, and the bioinformatics and computing service of Roscoff’s Biological Station (Abims platform) for computing time. This work was supported by the French Agence Nationale de la Recherche (project SEAD, ANR-13-ADAP-0011 and project SexChange, ANR-14-CE02-0001).

